# Feedback Control in Planarian Stem Cell Systems

**DOI:** 10.1101/029009

**Authors:** Marc Mangel, Michael B. Bonsall, Aziz Aboobaker

**Author notes:** Author for Correspondence: Marc Mangel.

## Abstract

**Background** In planarian flatworms, the mechanisms underlying the activity of collectively pluripotent adult stem cells (neoblasts) and their descendants can now be studied from the level of the individual gene to the entire animal. Flatworms maintain startling developmental plasticity and regenerative capacity in response to variable nutrient conditions or injury. We develop a model for cell dynamics in such animals, using methods of nonlinear dynamics, assuming that fully differentiated cells exert feedback control on neoblast activity. **Results** Our model predicts a number of whole organism level and general cell biological and behaviours, some of which have been empirically observed or inferred in planarians and others that have not. As previously observed empirically we find: 1) a curvilinear relationship between external food and planarian steady state size; 2) the fraction of neoblasts in the steady state is constant regardless of planarian size; 3) a burst of controlled apoptosis during regeneration after amputation as the number of differentiated cells are adjusted towards their correct homeostatic/steady state level. In addition our model describes the following properties that can inform and be tested by future experiments: 4) the strength of feedback control from differentiated cells to neoblasts (i.e. the activity of the signalling system) and from neoblasts on themselves in relation to absolute number depends upon the level of food in the environment; 5) planarians adjust size when food level reduces initially through increased apoptosis and then through a reduction in neoblast self-renewal activity; 6) following wounding or excision of differentiated cells, different time scales characterize both recovery of size and the two feedback functions; 7) the temporal pattern of feedback controls differs noticeably during recovery from a removal or neoblasts or a removal of differentiated cells; and 8) the signaling strength for apoptosis of differentiated cells depends upon both the absolute and relative deviations of the number of differentiated cells from their homeostatic level.

**Conclusions** We offer the first analytical framework for organizing experiments on planarian flatworm stem cell dynamics in a form that allows models to be compared with quantitative cell data based on underlying molecular mechanisms and thus facilitates the interplay between empirical studies and modeling. This framework is the foundation for studying cell migration during wound repair, the determination of homeostatic levels of differentiated cells by natural selection, and stochastic effects.

## Background

Stem cell systems operate by demand control [1-3] in which the needs of the organism determine in large part the behavior of the stem cells. Indeed, both cancer and ageing may be understood as failures of this feedback control, albeit in different ways. The highly regenerative planarian flatworms (Tricladdida), particularly *Dugesia* and *Phagocata* species, have been key models in the study of regeneration and wound healing for more than 100 years (see [4-6] for some classic studies; [7-11] for more recent ones). Their simplicity and the ease with which regeneration experiments can be performed make them an attractive system to understand the fundamental mechanics of regeneration. Recent advances in molecular techniques have allowed deeper understanding of these apparently simple organisms; for example, it is now possible to study the stem cell system and its descendants from the level of the single gene to the entire organism. The planarian life history provides the unique opportunity to take a systems approach to understanding stem cell dynamics in a whole organism.

In planaria, stem cells are called neoblasts and are defined collectively as the only dividing cells in the animal. Among these cells it has long been assumed that at least some cells are *bona-fide* pluripotent stem cells, capable of indefinite self-renewal and of producing all differentiated cell types in the adult animal; this was recently experimentally verified in the model species *Schmidtea mediterranea* [12]. A growing body of co-expression data shows that sub-populations of cycling neoblasts express lineage specifc mRNA markers [13]. Some of these co-expressed markers are functionally required for production of both the neoblast sub-population and the differentiated cell lineage in question; reviewed in [14]. This provides evidence for the existence of committed proliferating cells amongst the neoblast population, but this awaits definitive experimental proof.

Fully differentiated cells in planarians have been divided into about 15 different classes, or 3 super-classes (e.g. cells associated with metabolism, muscle, and the epidermis), with the actual number of functional cell types likely to be much higher [8,15]. Unlike other stem cell systems such as the bone marrow stem cell system, in planaria there is still no conclusive evidence for mitotically active progenitor cells with strictly limited potency [16-18]. There are however populations of transient post-mitotic stem cell progeny, and these cells either differentiate to a target lineage or potentially may apoptose rather than complete differentiation. We assume that the proportion of the various types of differentiated cells is regulated towards a homeostatic target [19, 20] but in this paper do not model how that target is set (see **Discussion**).

The requirement for a given mix of differentiated cells and the simplicity of the system make planarians an ideal system for studying homeostasis and regeneration, including scaling, reproductive fission, and responses to wounding and amputation [21, 22]. A minimum remaining tissue size is needed for such recovery after artificial amputation [23], but given that minimum size the entire organism can be regenerated through proliferation of stem cells to produce new tissue *de-novo* and remodeling of the remaining tissue [24].

We take a dynamical view of planarian stem cell system [25,26] and thus formulate our models as nonlinear dynamical systems in which the nonlinearity arises through feedback control. Although intraspecific variation exists, we ignore it for now.

We develop two models based on current knowledge of planarian stem cell biology. The first has neoblasts, non-mitotic progenitor cells, and three kinds of differentiated cells (three chosen for simplicity of setting parameters; the methods scale readily for an arbitrary number of differentiated cells). In the second model, we assume only one kind of differentiated cell and that the progenitor to differentiated cell transition is essentially instantaneous. This allows simplification that makes presenting some results clearer without losing any general principles.

We next give verbal and pictorial descriptions of the models, which are fully formulated in the **Methods** section. We then turn to **Results** beginning with steady state prediction of planarian size in relation to food in the environment and show that both the strength of feedback control in response to food in the environment and the constancy of the fraction of neoblasts are emergent properties of the model. We use the full model to explore growth, shrinkage, and regrowth under sufficient resources to maintain metabolism and to explore bursts of cell activity during remodeling following a fission or wounding. We use the simplified model to repeat the study of growth, shrinkage, and regrowth and then use it in two kinds of *in silico* experiments. In the first experiment, we ‘wound’ the planarian by removing a large number of differentiated cells, while in the second experiment we simulate death of neoblasts as happens after *γ* irradiation [27].

### Verbal and Pictoral Description of the Models

In the **Methods** we describe the models as a discrete time dynamical system (obtained by writing a system of differential equations in a form suitable for numerical solution). Here we give both verbal and pictorial (Figure 1) descriptions of the model. In Figure 1a we show the cell types and their transitions, along with the resource pool *Q*. We assume that neoblasts, *N*, are immortal and do not directly transition to differentiated cells, but rather go through a progeny *P* cell that is no longer mitotically active but which is not fully differentiated. Such a progeny cell may complete differentiation to one of the kinds of fully differentiated cells (in the figure, we show three kinds of such cells, indexed as *D_i_*) or may apoptose and thus return to the resource pool. Thus, the transitions for neoblasts are *N* → 2*N*; *N* → *N*, *P*; and *N* → *P, P*. The transitions for progeny cells are *P* → *Q* and *P* → *D*_1_, *P* → *D*_2_, or *P* → *D*_3_.

**Figure 1.**
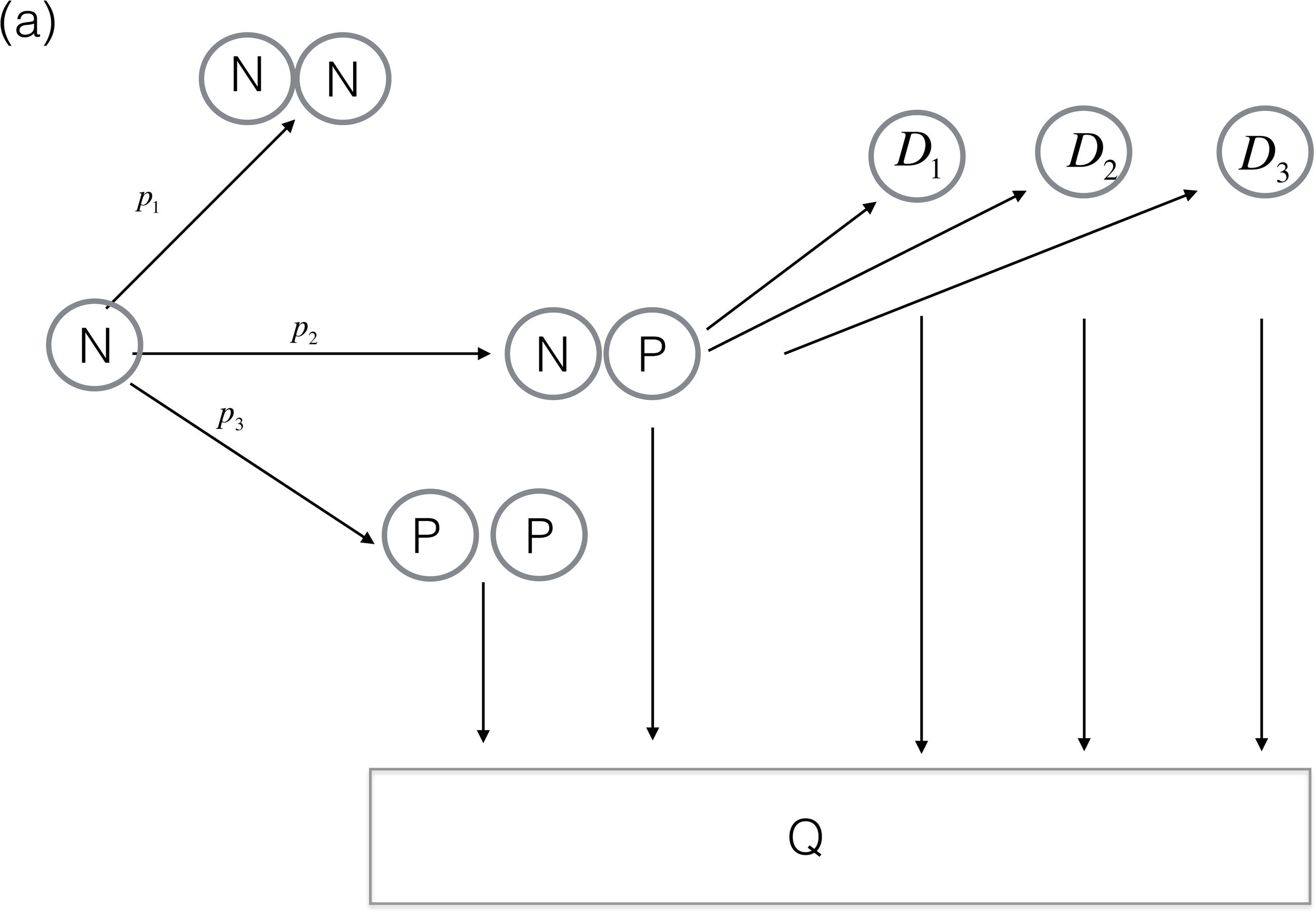
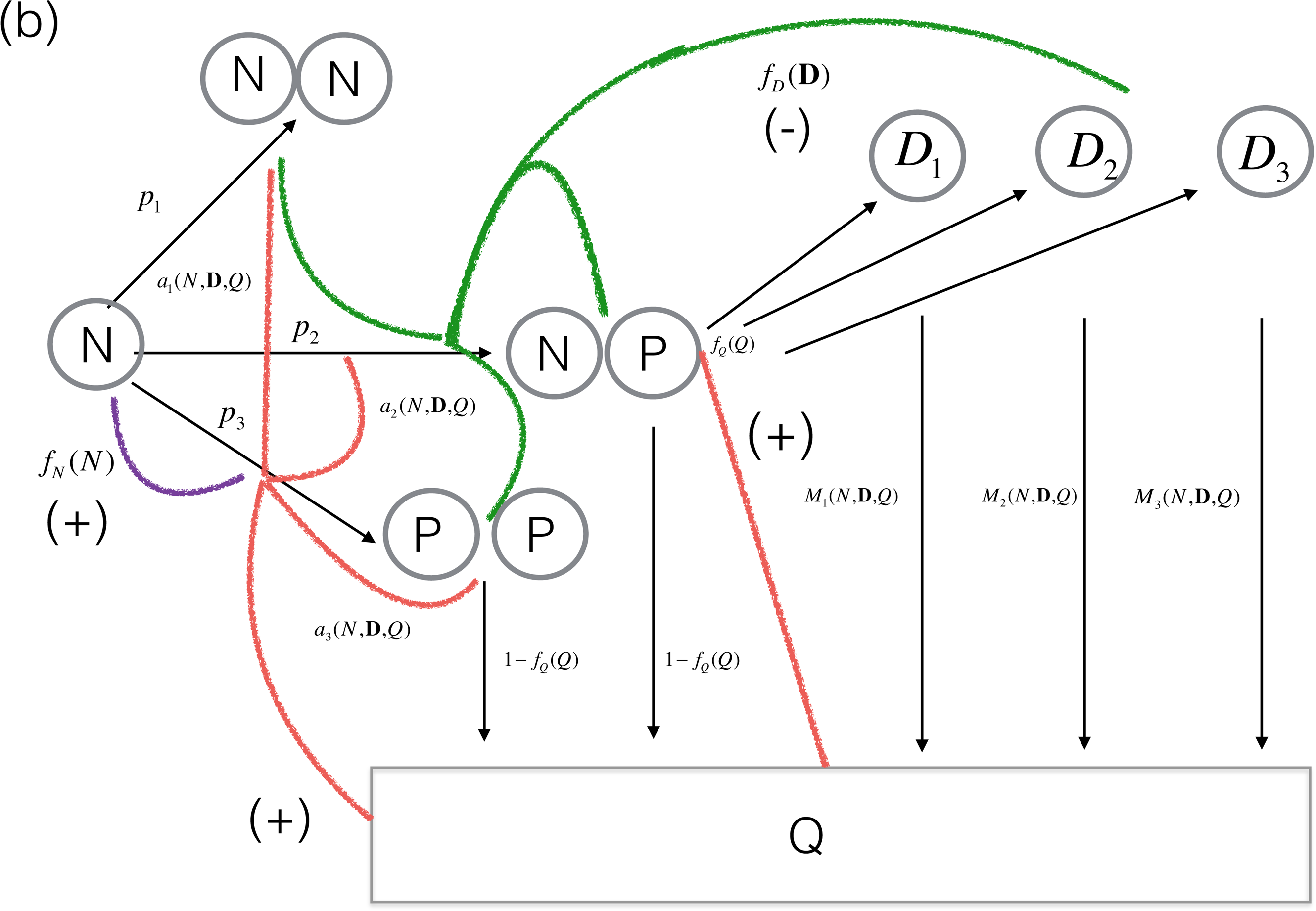
a) Pictorial description of the model involving neoblasts (*N*), non-mitotic progenitor cells (*P*), three classes of differentiated cells (*D_i_*), and a resource pool. The maximum rates of asymmetric renewal, symmetric differentiation, and asymmetric differentiation are *p*_1_, *p*_2_, and *p*_3_ respectively. As explained in the methods, a steady state exists only if *p*_1_ < *p*_3_. Progenitor cells either complete differentiation or return to the resource pool. In the absence of external injury, death fully differentiated cells apoptose and return to the resource pool. b) Feedback controls superimposed upon the transitions. These transitions have a maximum rate that is modified by feedback control. In Figure 1b, we superimpose feedback control links on these transitions. Beginning from the left of the figure, neoblasts exert positive feedback *f_N_*(*N*) (purple line) on the *N* → *P, P* transition in the sense that as the number of neoblasts increases, *f_N_*(*N*) increases, bounded by 0 and 1. The resource pool *Q* also has positive feedback on all transitions, in the sense that with larger resource pools the rate of transition is higher. However, we assume that the resource pool operates differentially on the different transitions, so use *a*_1_(*N*, **D**, *Q*), *a*_2_(*N*, **D**, *Q*), *a*_3_(*N*, **D**, *Q*) to indicate the feedback control of the resource pool on the transitions *N* → 2*N*, *N* → *N, P*, and *N* → *P, P* respectively. Finally, the differentiated cells exert negative feedback control on the transitions, in the sense that as the number of differentiated cells increases, the rates of transitions decline, sharing a common feedback control *f_D_* (**D**). Here we assume the that the differentiated cell in shortest supply sets the feedback control. Absent an external source of mortality, the only transition for differentiated cells is *D*_1_ → *Q* through cell death, which occurs for cell type *i* at rate *M_i_*(*N*, **D**, *Q*) where **D** is the vector (*D*_1_, *D*_2_, *D*_3_). In addition, progenitor cells may either fully differentiate or return to the resource pool through apoptosis. We assume that the rate of the former is determined by a function *f_Q_*(*Q*) that increases as the size of the resource pool increases.

These transitions have a maximum rate that is modified by feedback control. In Figure 1b, we superimpose feedback control links on these transitions. Beginning from the left of the figure, neoblasts exert positive feedback *f_N_* (*N*) (purple line) on the *N* → *P, P* transition in the sense that as the number of neoblasts increases, *f_N_*(*N*) increases; it ranges between 0 and 1. The resource pool *Q* also has positive feedback on all transitions, in the sense that with larger resource pools the rate of transition is higher. However, we assume that the resource pool operates differentially on the different transitions, so use *α*_1_(*N*, D, *Q*), *a*_2_(*N*, D, *Q*), *a*_3_(*N*, D, *Q*) to indicate the feedback control of the resource pool on *N* → 2*N*, *N* → *N*, *P* and *N* → *P*, *P* respectively. Finally, the differentiated cells exert negative feedback control on the transitions, in the sense that as the number of differentiated cells increases, the rates of transitions decline, sharing a common feedback control *f_D_*(**D**) where **D** is the vector (*D*_1_, *D*_2_, *D*_3_). We assume the that the differentiated cell in shortest supply sets the feedback control (details given in **Methods**).

In the absence of an external source of mortality, the only transition for differentiated cells is *D_i_* → *Q* through cell death, which occurs for cell type *i* at rate *M_i_*(*N*, D, *Q*). In addition, progenitor cells may either fully differentiate or return to the resource pool through apoptosis. We assume that the rate of the former is determined by a function *f_Q_*(*Q*) that increases as the size of the resource pool increases.

In Figure 2, we show the feedback functions and the resource dependent rate of apoptosis. The feedback function *f_D_* (*D*) (Figure 2a) from differentiated cells on the transitions of neoblasts (green line in Figure 1b) falls from 1.0 as the number of differentiated cells increases. This function has a parameter *α* that controls the rate of decline, in the sense that larger values of *α* mean smaller values of *f_D_* (*D*) at the same level of differentiated cells. The feedback control from neoblasts to asymmetric differentiation (purple line in Figure 1b, Figure 2b) is a sigmoidal function that increases towards 1.0 as the number of neoblasts (measured as the fraction of the steady state value) increases. It is characterized by two parameters: the value of neoblasts at which *f_N_*(*N*) = 0.5 and the spread around that value. The feedback functions *a_i_*(*N*, D, *Q*) and the resource dependent rate of mortality of differentiated cells depend, for a model with three kinds of differentiated cells, on five variables. In Figure 2c, we show a cross-section of those functions by holding the number of neoblasts and differentiated cells constant and only varying the resource level. The values shown here are illustrative of the shape and relationship of the three feedback functions. The x-axis intentionally has no units since these images are intended to be schematic rather than accurate representations. In Figure 2d we show the resource dependent rate of natural mortality, again in cross-section. Full details are in the **Methods**.

**Figure 2.**
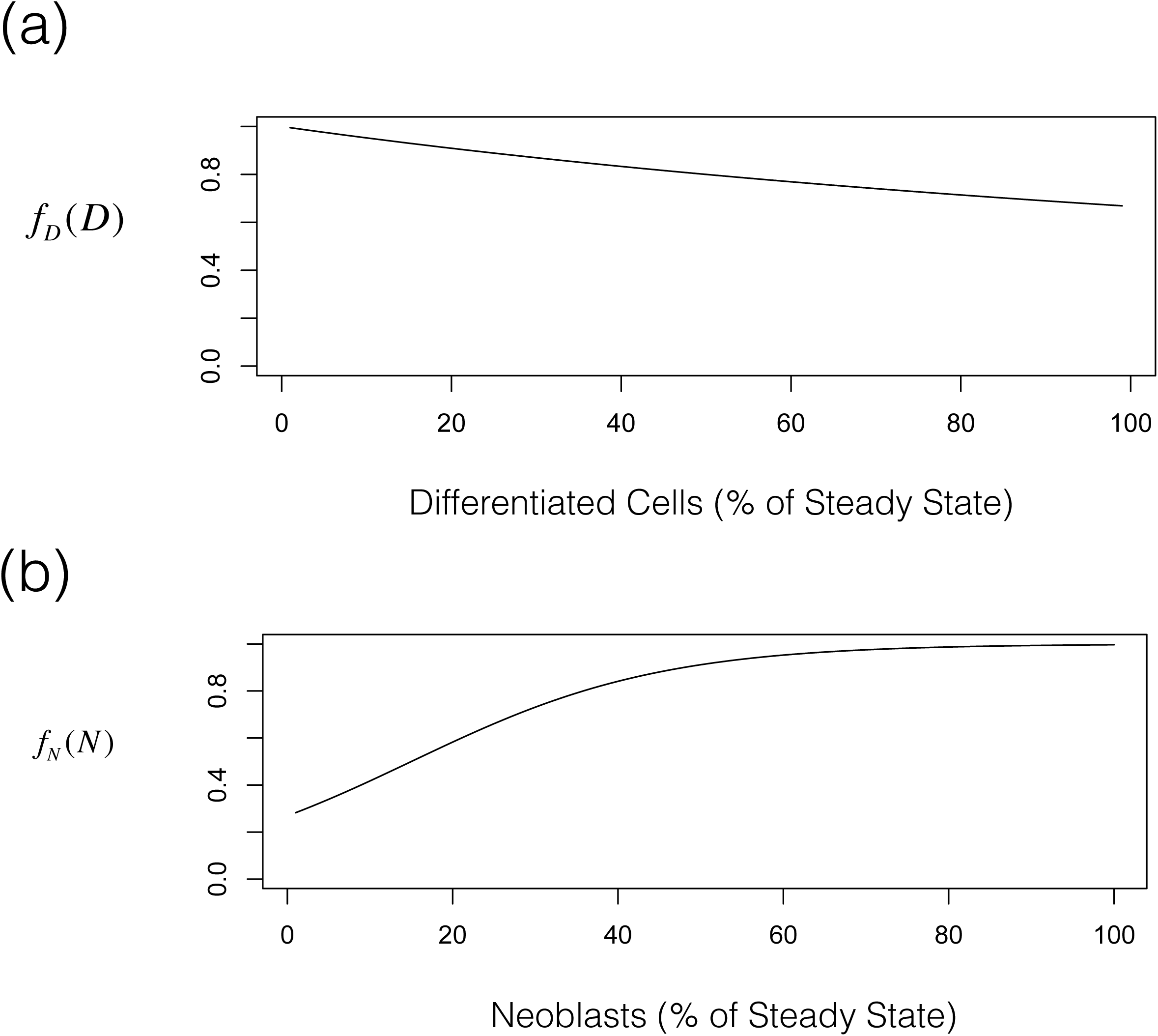
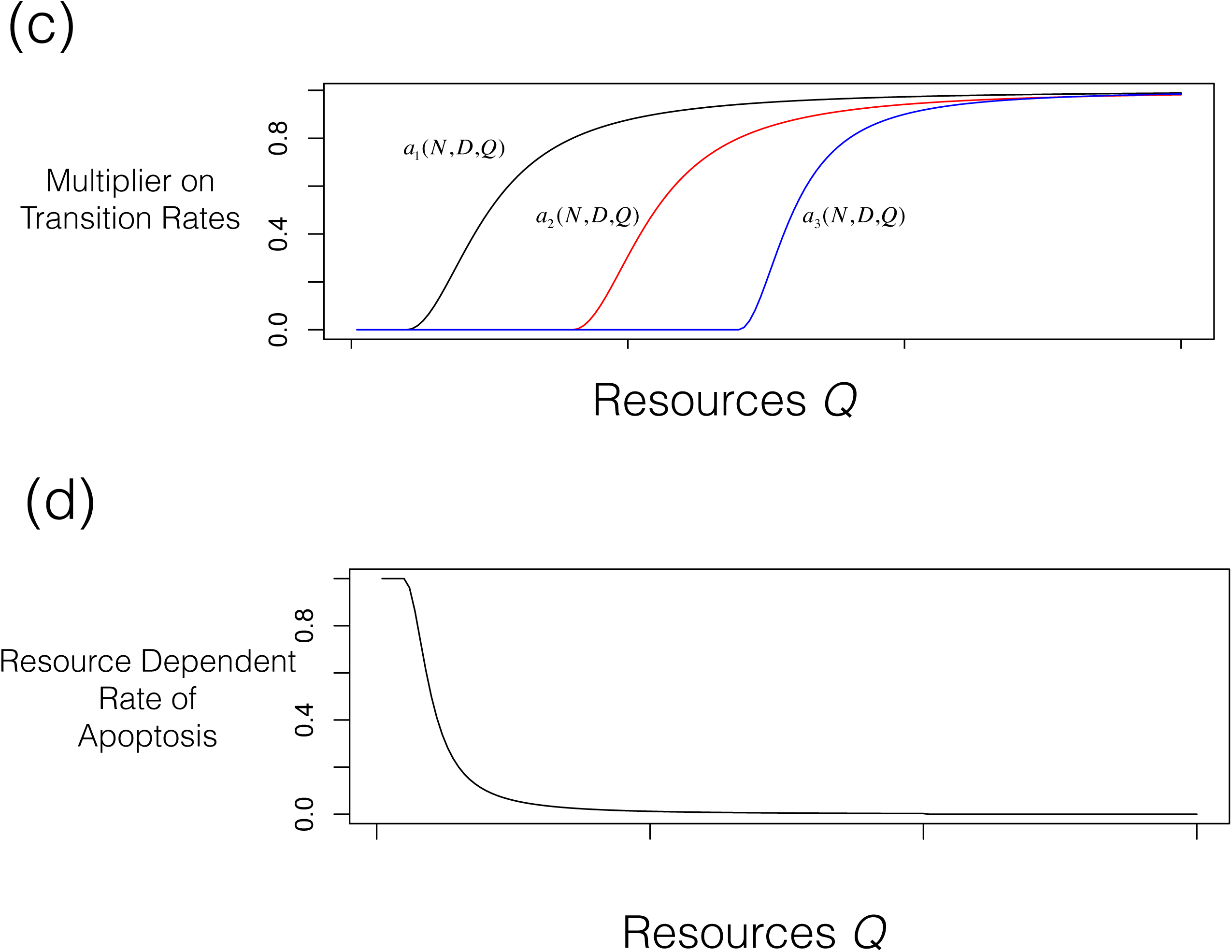
a) The feedback function *f_D_* (*D*) from differentiated cells on the transitions of neoblasts (green line in Figure 1b) falls from 1.0 as the number of differentiated cells (here represented as the fraction of the steady state value) increases. b) The feedback control from neoblasts to asymmetric differentiation (purple line in Figure 1b) increases towards 1.0 as the number of neoblasts (here represented as the fraction of the steady state value) increases. c) The feedback functions *a_i_*(*N*, **D**, *Q*) depend, for a model with three kinds of differentiated cells, on five variables. Here we show a ‘cross-section’ of those functions by holding the number of neoblasts and differentiated cells constant and only varying the resource level. All of these are arbitrary but the shape and relationship of the three feedback functions are not. d) Similarly, the resource dependent rate of mortality of differentiated cells depends on the number of neoblasts and differentiated cells through their metabolic requirements and here we show a cross-section holding those cell numbers constant and varying resource levels. The x-axis intentionally has no units since these images are intended to be schematic rather than accurate representations. Full details are in the **Methods**.

To capture the transition from progenitor to fully differentiated cell, we make the *relative need* assumption that progenitors transition to differentiated cells according to how far they deviate from the homeostatic level, which is determined by both the number of differentiated cells and their relative proportion. The rate of apoptosis of differentiated cells is also determined by the number of differentiated cells and how far they deviate from the homeostatic proportion, and the resource pool. In particular, the rate of mortality increases for numbers of, or proportions of, differentiated cells above the homeostatic level and also increases as the resource pool declines.

In the simplified model, we compress all of the differentiated cells into a single type and assume that the progenitor transitions are so rapid that they can be ignored. This allows clearer analytical and pictorial representation of the feedback controls. Details are given in **Methods**.

## Results and discussion

### Overview

We begin with steady state results, using the full model, showing the relationship between food in the environment and planarian size, and how the strength of feedback control from differentiated cells to neoblast activity emerges in response to food in the environment. We also demonstrate that in the steady state the fraction of neoblasts is independent of size (compare with with empirical work in [28]). We then use the full model to study growth, shrinkage, and regrowth with sufficient resources. When resources are ample, feedback control is independent of resources and only depends on the number of differentiated cells (i.e., the *a_i_*(*N*, D, *Q*) = 1 and *f_Q_*(*Q*) = 1). We then turn to remodeling of a planarian following fission. To do, this we assume that the initial cell numbers are a small fraction of their steady state values and that neoblasts and differentiated cells are out of their homeostatic levels. Thus we anticipate that as the planarian regenerates it will need to change the number of neoblasts and mixture of differentiated cells (decreasing some that remain in excess) so that there will be a burst of mortality following fission, increasing the resource pool which is then used to regrow towards the steady state. We use the simplified model to study cell activities during growth, shrinkage, regrowth, and regeneration, and in the *in-silico* experiments in which a fraction of the differentiated cells are removed, as would happen with wounding or amputation, or a fraction of the neoblasts are removed, as would happen with a relatively precise X-ray treatment at the center of the planarian.

### Steady States

In Figure 3 we show the steady state size of the planarian for the full model; this is determined by the steady state number of cells of all types and an allometric relationship between size and total cell number (data in [29]; Eqn 26 below). The curvilinear nature of this relationship is due to not all cells being able to acquire resources from the external environment (Eqn. 12 below).

**Figure 3.**
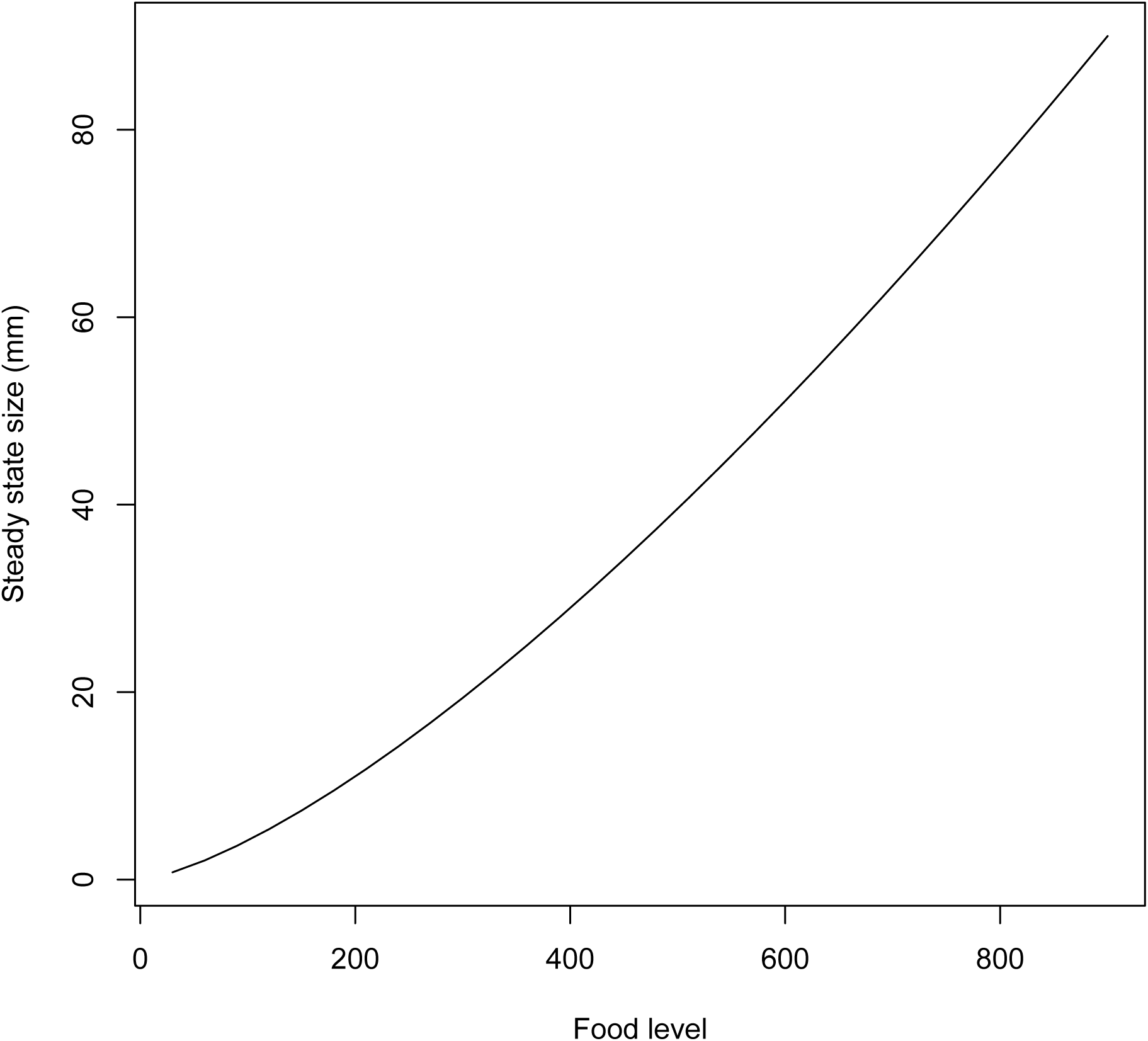
The steady state population size for the full model as a function of food in the environment (based on equations 21–26).

As described in detail in the methods, the strength feedback control of is an emergent property, depending upon the level of food in the environment and the maximum rates of asymmetric renewal and differentiation (Figure 4). The strength of feedback control from differentiated cells to neoblast activity declines as food in the environment increases because the increasing food supports more cells overall. Thus, we predict that the strength of the signaling system between differentiated cells and neoblast activity will respond to external food.

**Figure 4.**
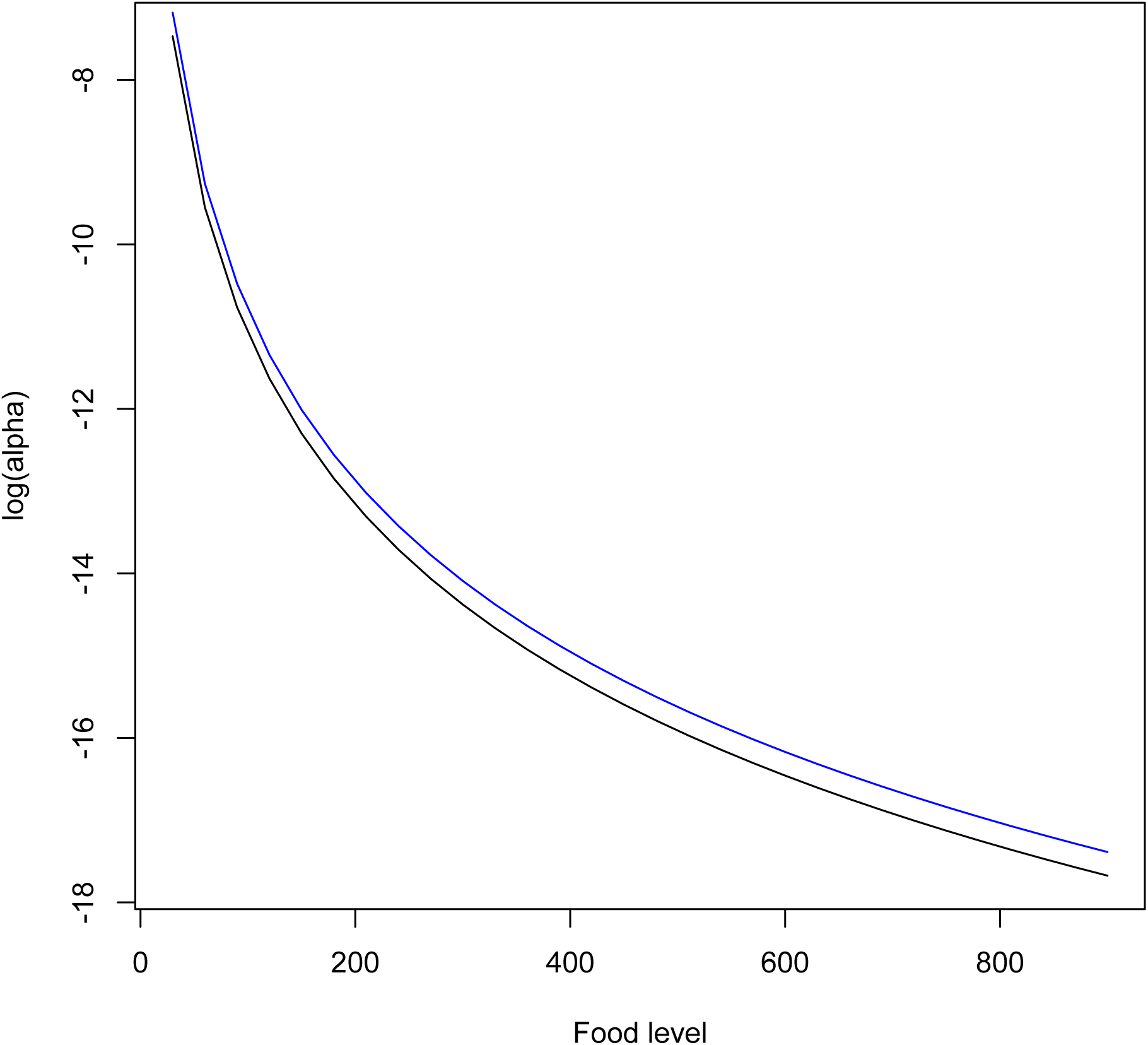
The strength of feedback control from differentiated cells to neoblasts emerges as food-dependent from Eqn 2. Here we show two curves: one for the differentiated cells with slightly lower rate of natural mortality and one for the two kinds of differentiated cells with slightly higher rate of natural mortality.

In Eqns 15-21, we show that the fraction of neoblasts in the steady state is constant, independent of the size of the planarian. This proportion is determined by the transition rates and rates of death of differentiated cells.

### Dynamics with Sufficient Food Resources (Full Model)

In Figure 5, we show the temporal pattern of size, total mortality of differentiated cells, total mortality of differentiated cells, and fraction of neoblasts under growth, shrinkage, and regrowth for a situation in which the planarian starts at 20% of its steady state size in an environment with food availability *Y_e_* = 450. Shrinkage can occur with sufficient resources due to apoptosis of differentiated cells that return to the resource pool. At time 70,000 food is dropped to *Y_e_* = 400 and then at time 140,000 increased to *Y_e_* = 480. Even in the absence of a resource constraint, the cell population dynamics show a dependence upon the level of food in the environment. This is due to the feedback control on neoblast activity from differentiated cells and feedback control on neoblasts themselves on asymmetric division.

**Figure 5.**
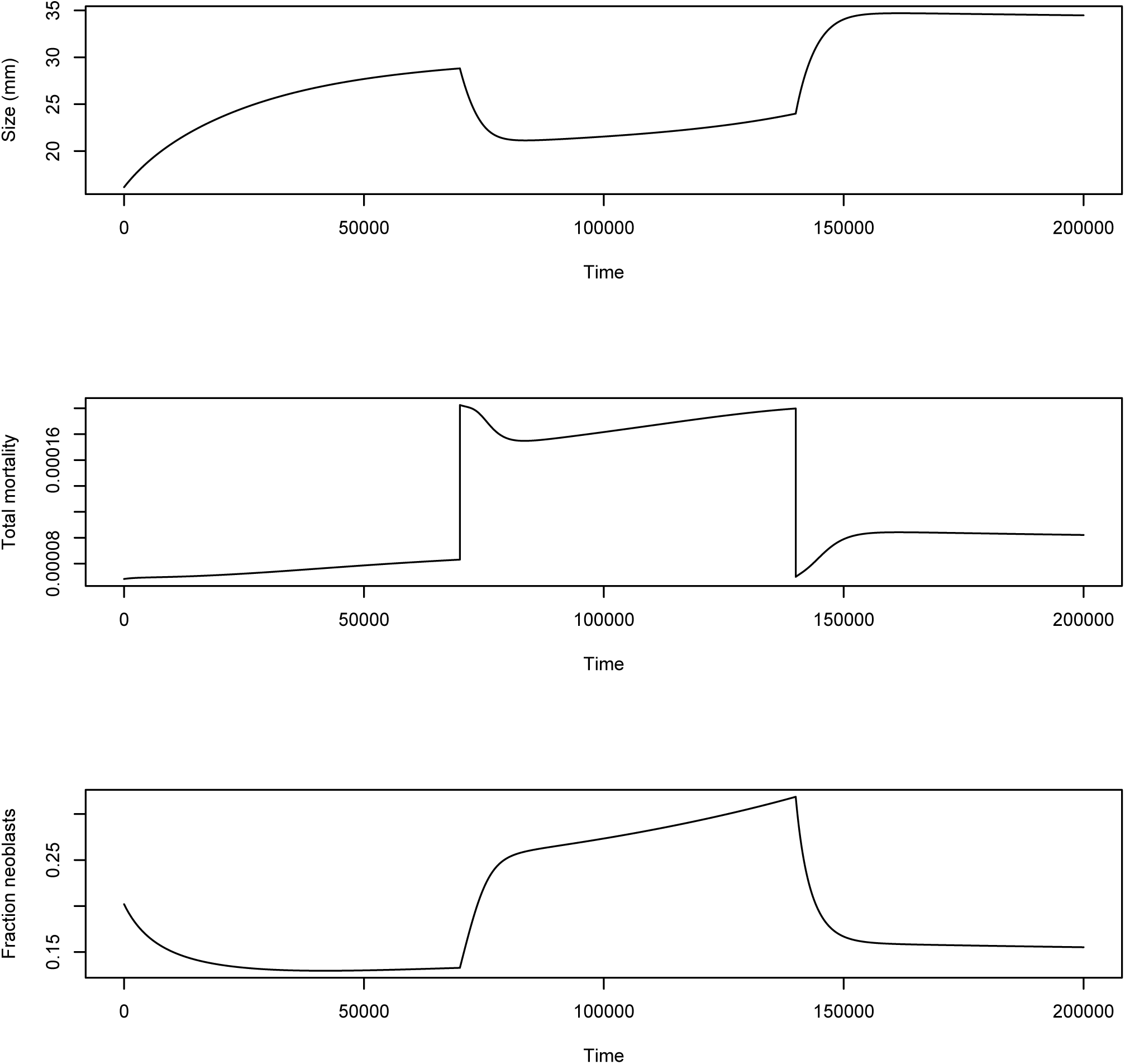
The temporal pattern of size, total mortality of differentiated cells, total mortality of differentiated cells, and fraction of neoblasts under growth, degrowth, and regrowth for a situation in which the planarian starts in an environment with *Y_e_* = 450 at 20% of its steady state size. At time 70,000 food is dropped to *Y_e_* = 400 and then at time 140,000 increased to *Y_e_* = 480. Even in the absence of a resource constraint, the cell population dynamics show a dependence upon the level of food in the environment. This is due to the feedback control on the activity of neoblasts.

### Remodeling Following a Fission (Full Model)

After a natural fission, the neoblast proportion is probably slightly higher than the normal steady state, but many differentiated cell types will be much lower than steady state and others will be higher. For example, in tail fragments neurons will be lower than the steady state and gut cells higher. In Figure 6, we show cell dynamics during remodeling following a fission. We assume that the initial cell numbers are 50% of the steady state number of neoblasts, and 10%, 30%, and 5% of the steady state values of the three kinds of differentiated cells respectively (rather than 25% neoblasts and 40%, 30%, 30% relative distribution of differentiated cells in the steady state). Starting from relatively small size, the planarian grows to a steady state (since food is constant) (upper panel), and a burst of mortality occurs (second panel) as the differentiated cells apoptose in order to achieve their target values (bottom panel). These results could be compared with empirical work in [30] in more detail but show qualitatively similar patterns.

**Figure 6.**
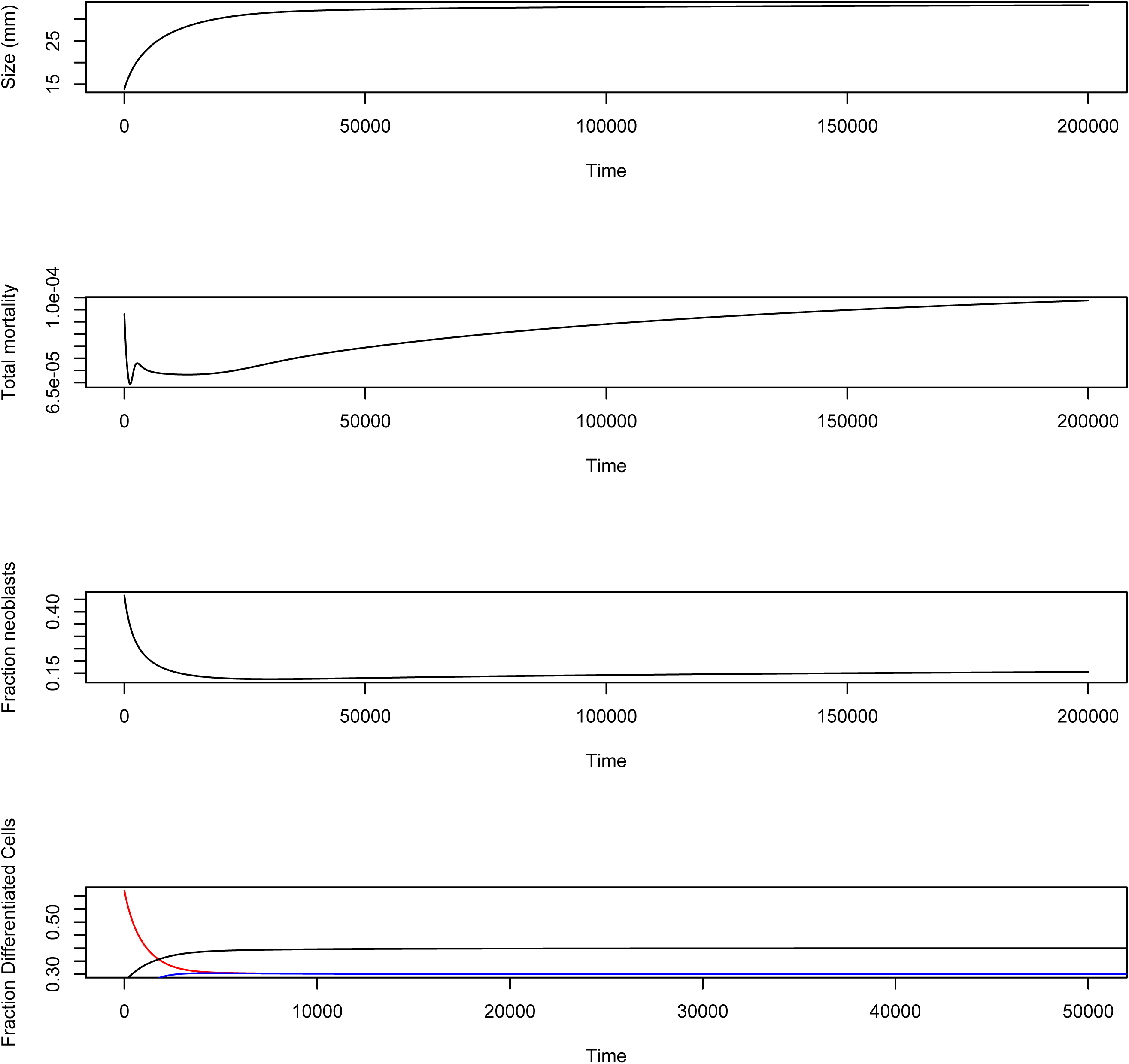
Celll dynamics during remodeling following a division. We assume that the initial cell numbers are 5% of the steady state number of neoblasts, and 10%, 30%, and 5% of the steady state values of the three kinds of differentiated cells respectively (rather than 25% neoblasts and 40%, 30%, 30% relative distribution of differentiated cells in the steady state). Starting from relatively small size, the planarian grows to a steady state (since food is constant) (upper panel), a burst of mortality initially (second panel) as the differentiated cells apoptose in order to achieve their target values (bottom panel), and a complicated trajectory for the fraction of neoblasts as the animal readjusts its composition.

### *In-silico* Experiments (Simplified Model)

We first repeat growth, shrinkage, and regrowth using the simplified model (Figure 7) to verify the same qualitative patterns. We started the planarian at 20% of the steady state size associated with food level *Y_e_* = 240 and then varied food over time as shown in Figure 7a. The planarian grows towards its new steady state until food is decreased at *t* = 70, 000 and then again at *t* = 150, 000 and in response the planarian size decreases (Figure 7b). During this entire process, however, the fraction of neoblasts is nearly constant (Figure 7c). The feedback controls *f_D_*(*D*(*t*) and *f_N_*(*N*(*t*)) (Figures 7d and 7e respectively) respond to the food pattern in complex ways, in part because *α* in the feedback control 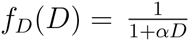 is a function of external food level (Figure 4) and because of additional mortality when resources are insufficient. On the other hand the resource dependence of the three neoblast transitions (Figure 7f) – measured by (i.e. *p*_1_*a*_1_ (*N, D, Q*), *p*_2_*a*_2_(*N, D, Q*) *f_D_*(*D*), and *p*_3_*a*_3_(*N, D, Q*) *f_D_*(*D*)*f_N_*(*N*) showed much simpler patterns. Indeed it is only the *N* → *N, P* transition that shows a response to the availability of food. Whether the relative insensitivity of the *a_i_*(*N, D, Q*) is a general property or due to the formulation or the specific parameter values is still not clear but a full analysis is beyond the scope of this paper.

**Figure 7.**
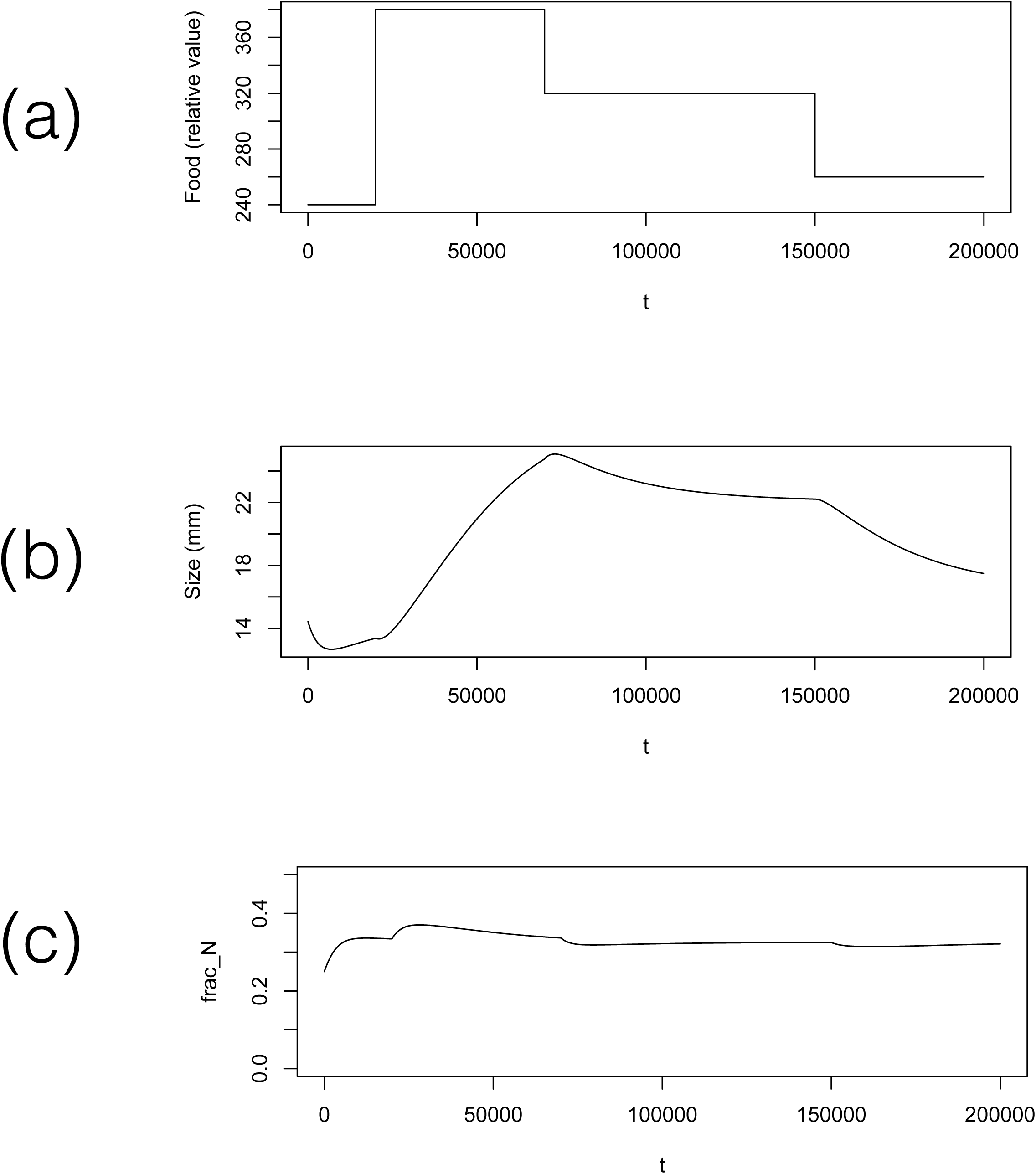
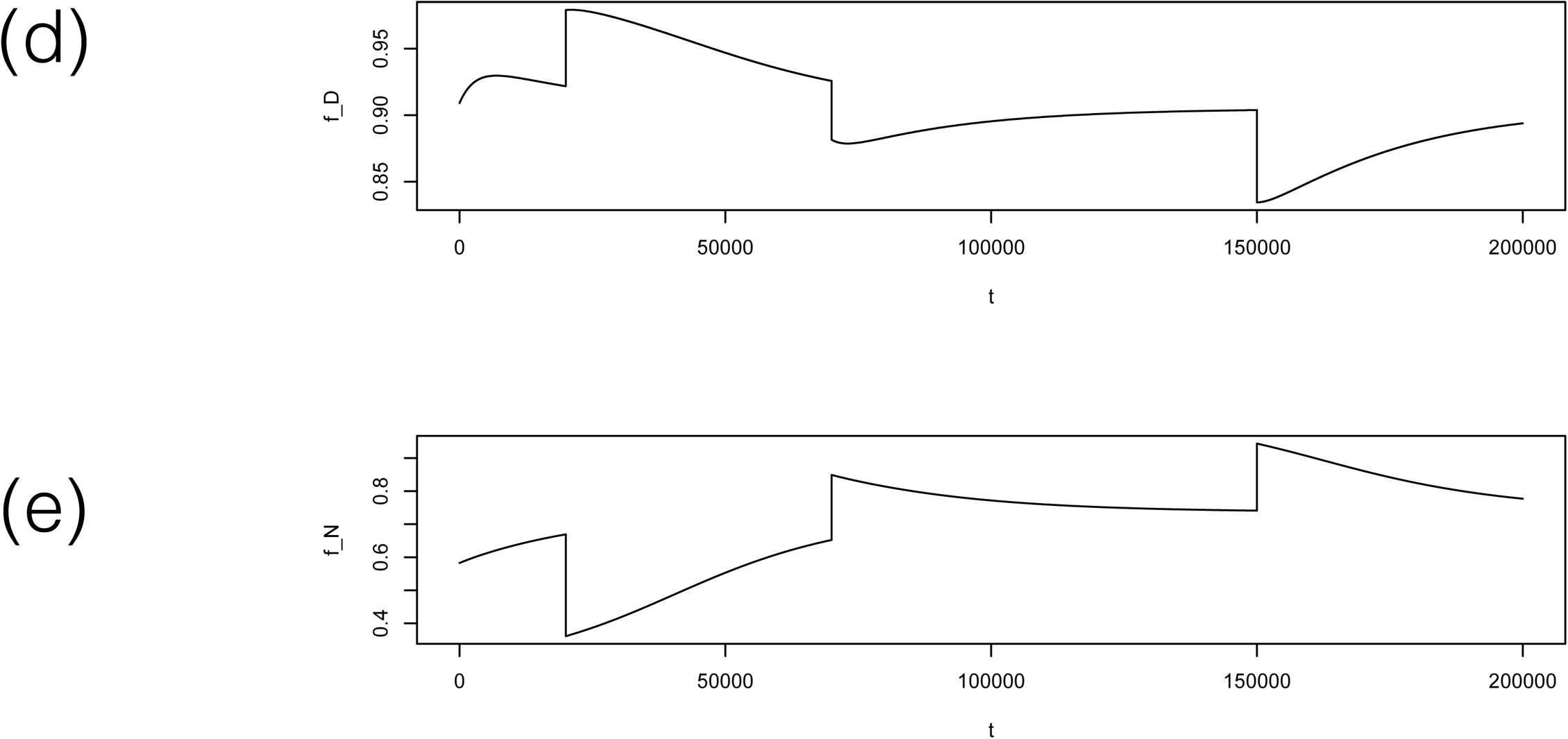
Pattern of growth and shrinkage using the simplified model. a) The temporal pattern of food. b) Predicted size of the planarian as a function time. c) The fraction of neoblasts as a function of time. d) The feedback control from differentiated cells to neoblast activity. e) The feedback control from neoblasts to themselves on the *N* → *P, P* transitions. e) The activity of neoblasts (i.e. *p*_1_*a*_1_(*N, D, Q*), *p*_2_*a*_2_(*N, D, Q*)*f_D_*(*D*), and *p*_3_*a*_3_(*N, D, Q*)*f_D_*(*D*)*f_N_*(*N*) for asymmetric renewal, symmetric renewal, asymmetric division respectively (upper to lower panels).

In Figure 8, we show the results of the *in silico* excision experiment in which we allow the planarian to grow to its steady state size for the given food environment and then at *t* = 70,000 remove differentiated cells via ‘excision’. This leads to an increase in the fraction of neoblasts (Figure 8b), an increase in the feedback control function *f_D_* (*D*) (Figure 8c) and a slight drop in *f_N_* (*N*) (Figure 8d, note the vertical scale. Also note the different time scales in the recovery of size and the two feedback functions. Size is recovered by about t=85,000 and although the feedback function *f_D_* returns to its previous value at approximately the same time, the feedback function *f_N_* takes much longer to return to its steady state. These results can be compared to experiments in [31].

**Figure 8.**
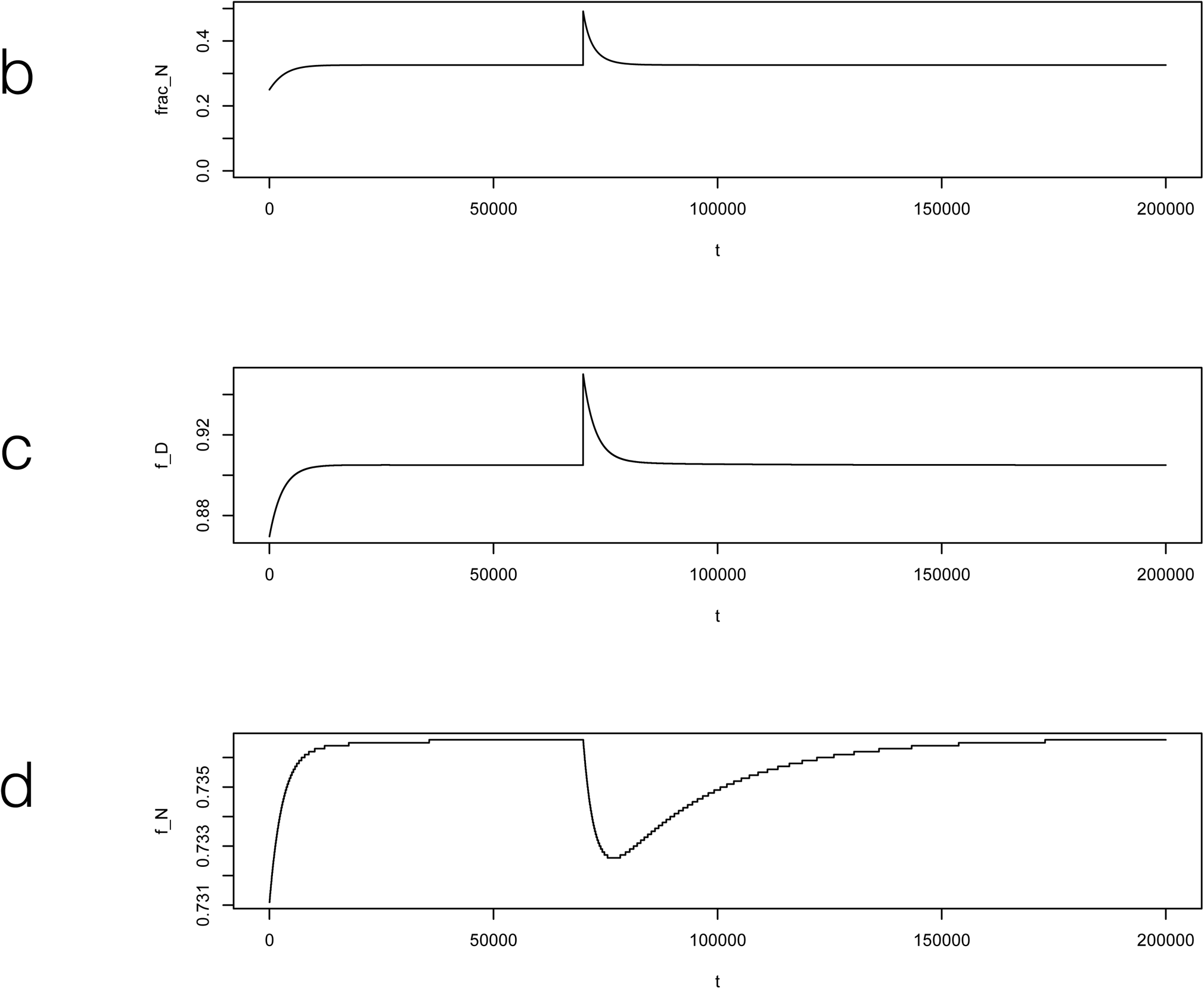
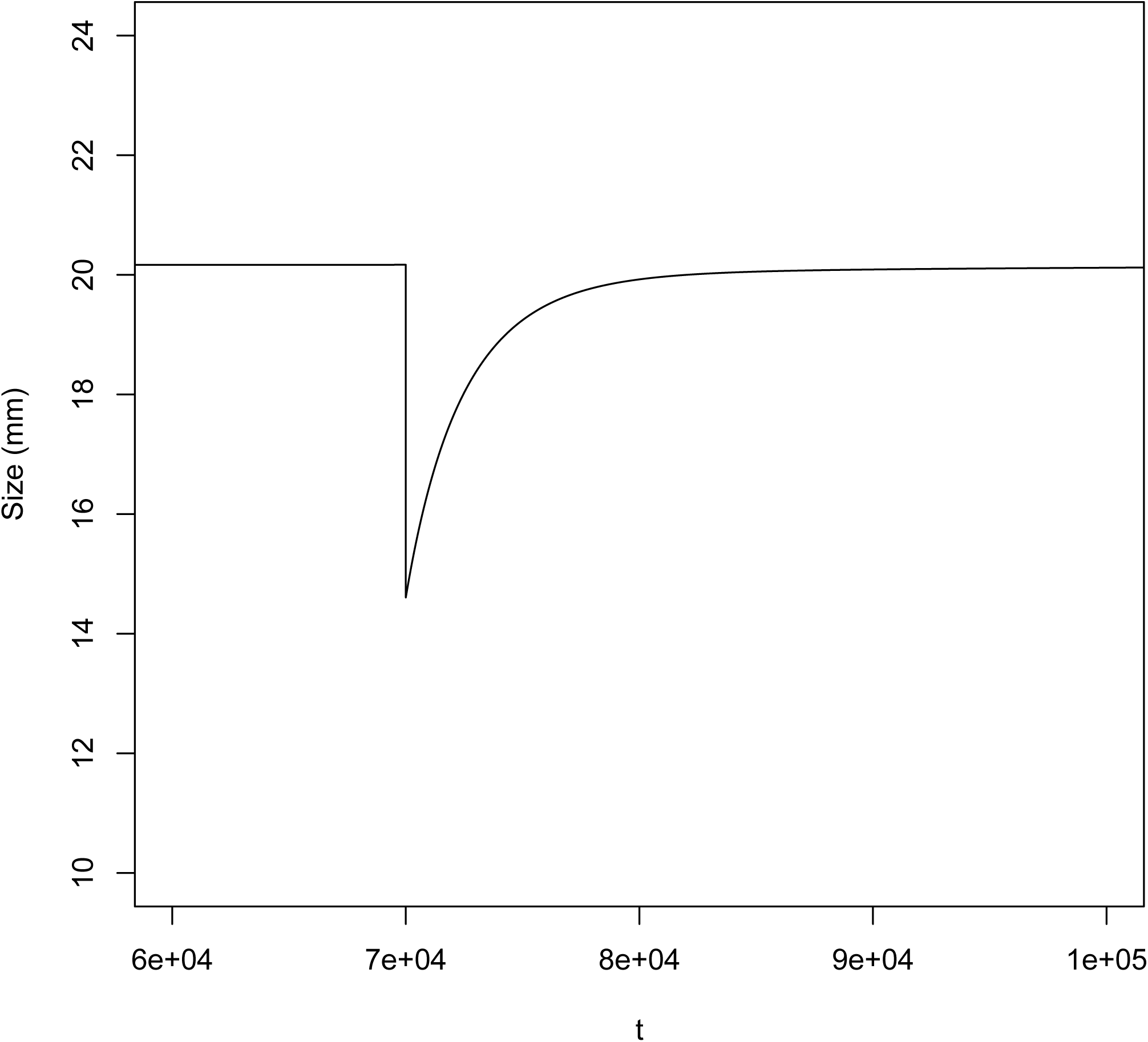
The *in silico* excision experiment. a) After the planarian has reached its steady state size at *t* = 70, 000 differentiated cells are removed by excision, leading to a drop in size. b) This leads to an increase in the fraction of neoblasts (upper panel), an increase in the feedback control function *f_D_* (*D*) (middle panel) and a slight drop in *f_N_* (*N*) (lower panel, note the vertical scale.

In Figure 9 we show the results of a similar ‘X-ray’ experiment: After the planarian has reached its steady state size at *t* = 70, 000 neoblasts are removed. Since we assume that size is determined by differentiated cells only there is no change in size, but an increase in the feedback control function *f_D_*(*D*) (middle panel), and a significant drop but then recovery of *f_N_*(*N*). Notice the very different patterns of the feedback control functions in Figures 8 and 9, suggesting that we can predict the pattern of cell-signalling systems based on the *in silico* experiments.

**Figure 9.**
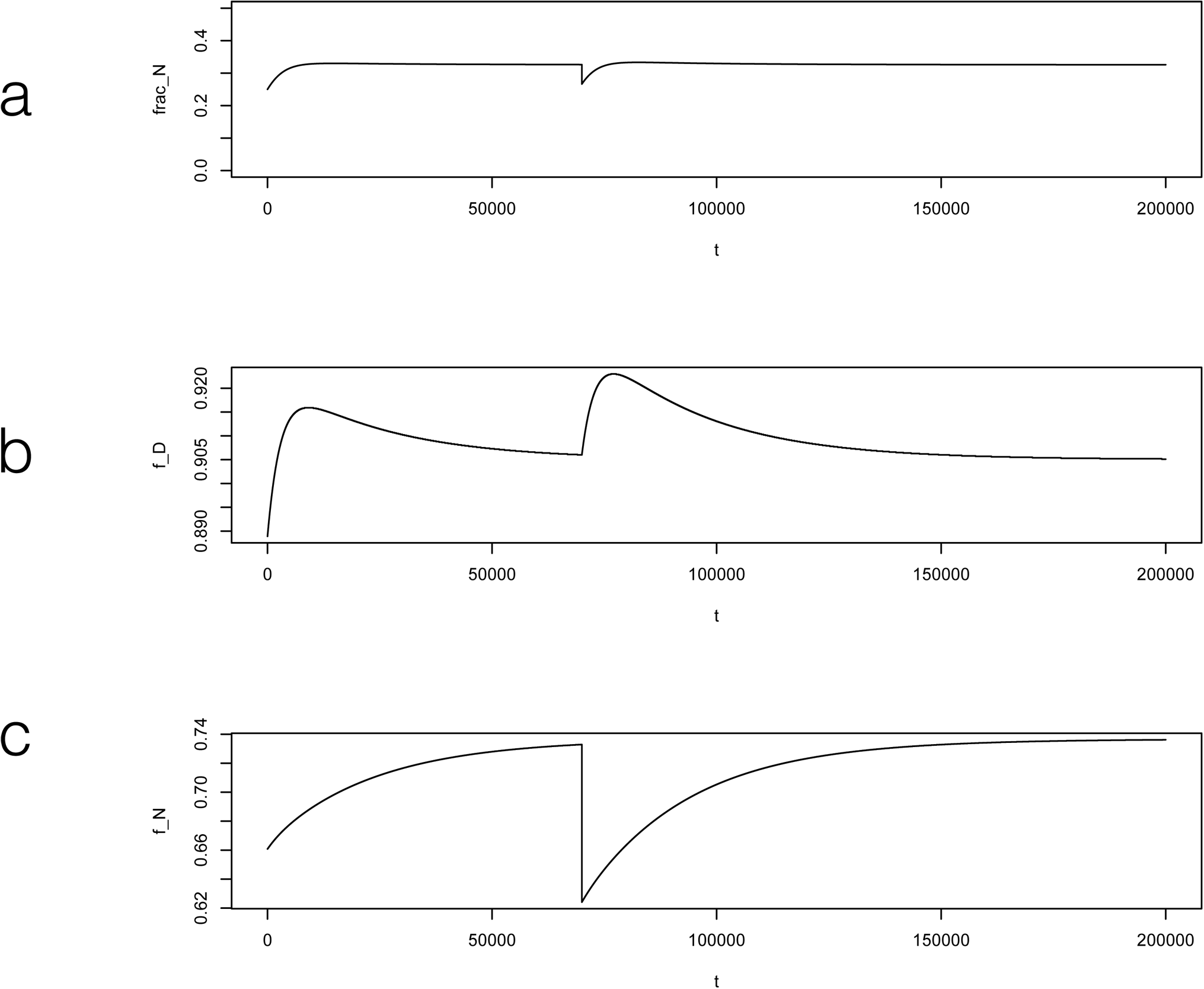
The *in silico* X-ray experiment. After the planarian has reached its steady state size at *t* = 70, 000 neoblasts are removed. This leads to a decrease in the fraction of neoblasts (upper panel), an increase in the feedback control function *f_D_*(*D*) (middle panel), and a significant drop but then recovery of *f_N_*(*N*) (lower panel).

## Conclusions

Our models lead to testable predictions about the dynamics of size, the rates of mortality, and the signaling systems in planaria.

These include:

1. a curvilinear relationship between external food and planarian steady state size;
2. that the strength of feedback control from differentiated cells to neoblasts (i.e. the activity of the signaling system) and from neoblasts on themselves depends upon the level of food in the environment;
3. the fraction of neoblasts in the steady state is constant regardless of planarian size;
4. planarians adjust size when food shifts first due to apoptosis and then through a reduction in neoblast activity;
5. a burst of mortality during regeneration as the number of differentiated cells are adjusted towards their homeostatic level;
6. following wounding or excision of differentiated cells different time scales characterize the recovery of size and the two feedback functions;
7. the temporal pattern of feedback controls differs noticeably during recovery from a removal of neoblasts or a removal of differentiated cells; and
8. the signalling for apoptosis of differentiated cells depends upon both the absolute and relative deviations of differentiated cells from their homeostatic level.

Much remains to be done, including comparing models to data explicitly, evolutionary origins of the steady state fraction [32], spatial dynamics of cells [33], and extension to stochastic models [34]. All of this requires the work here, which can thus be regarded as a first step towards the full modeling of the planarian stem cell system.

## Methods

We first describe the states that characterise the planarian, feedback control, and the dynamics of states. After that we determine the steady state and then discuss dynamics, and how *in silico* experiments can be performed to match empirical studies.

### States and Transitions

We let *N*(*t*) denote the number of neoblasts at time *t*, *D_i_*(*t*) the number of differentiated cells of type *i* at time *t*, for *i* = 1, 2,&*I* (in computations we set *I* = 3), and *D*(*t*) = (*D*_1_(*t*), *D*_2_(*t*), *&D_I_*(*t*)) the entire collection of differentiated cells.

We denote the number of non-mitotic but undifferentiated progeny and resource pool at time *t* by *P*(*t*) and *Q*(*t*) respectively. We let lower case indicate specifc values of these dynamic variables. We consider three kinds of transitions: 1) asymmetric neoblast renewal, i.e. *N* → 2*N*; 2) symmetric renewal and progeny production, i.e. *N* → *N* + *P*; and 3) asymmetric progeny production, i.e. *N* → 2*P*. A progeny cell either returns to the resource pool or continues to complete differentiation into one of the types of differentiated cells, as explained below.

### Feedback Control

Each of the transitions 1-3 above are subject to feedback control [3, 26, 34]. Suppose that *N*(*t*) = *n*, **D**(*t*) = **d** and *Q*(*t*) = *q*.

We assume that the fraction of neoblasts undergoing the *N* → 2*N* transition is *p*_1_*a*_1_ (*n, q*) where *p*_1_ is a fixed value (we explain below how it is set, in the section on parameters) and *a*_1_ (*n*, **d**, *q*) is determined as follows. If *m_r_* is the metabolic rate of neoblasts and differentiated cells (which we assume, for simplicity, to be the same) and *m_d_* is the cost of division, then 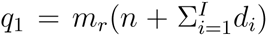 is the metabolic cost of maintenance of the existing cells and *q*_2_ = *q*_1_ + *m_d_p*_1_*n* is the level of resources needed to maintain all existing cells and support all asymmetric neoblast renewals. We let *q*_12_ denote the average of *q*_1_ and *q*_2_ and model the feedback control on *N* → 2*N* divisions by a Hill-type function

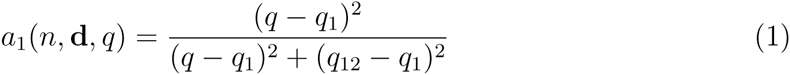

as long as *q* > *q*_1_; otherwise we set *a*_1_(*n*, **d**, *q*) = 0.

The transitions *N* → *N* + *P* and *N* → 2*P* involve resource-dependent and cell-number dependent feedback control. We assume that the fraction of neoblasts undergoing *N* → *N* + *P* is *p*_2_*a*_2_(*n*, **d**, *q*)*f_D_*(**d**) where *p*_2_ is a fixed value, *a*_2_(*n*, **d**, *q*) is determined in a manner similar to above and

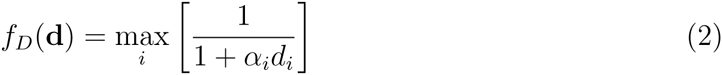

where *α_i_* is sets the strength of control from differentiated cells to neoblasts. The “max*_i_*” means that the differentiated cells that are in most demand set the level of feedback control. Given this control, *q*_3_ = *q*_2_ + *f_D_*(**d**)*m_d_n* is the level of resources needed to support all *N* → *N* + *P* transitions. We let *q*_23_ denote the midpoint of *q*_2_ and *q*_3_ and set

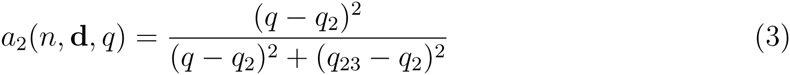

as long as *q > q*_2_; otherwise we set *a*_2_(*n*, **d**, *q*) = 0.

We assume that *N* → 2*P* involves an additional feedback control from neoblasts; so that when neoblast numbers are low this transition is suppressed. We set

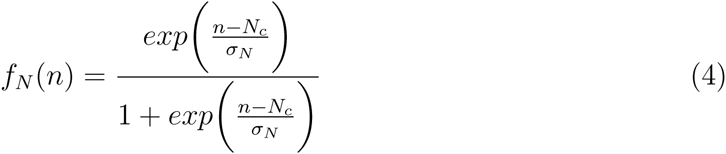

Since the exponent will be 0 when *n* = *N_c_*, *f_N_*(*N_c_*) = 0.5. The parameter *σ_N_* controls the sigmoidal or S-shape of *f_N_*(*n*). As *σ_N_* declines, *f_N_*(*n*) becomes more and more knife-edged, close to 0 when *n* < *N_c_*, close to 1 when *n* > *N_c_* but still 0.5 when equality holds. In the limit that *σ_N_* is very large (i.e. many times greater than *n* could be) *f_N_* (*n*) is close to 0.5 regardless of the value of *n*. We assume that *σ_N_* and *N_c_* are proportional to the steady state number of neoblasts, and are thus also environmentally determined by the level of food (see below, **Parameters**).

The fraction of neoblasts undergoing *N* → 2*P* transitions is *p*_3_*a*_3_(*n*, **d**, *q*) *f_D_*(**d**)*f_N_*(*n*) where *p*_3_ is a fixed value, and *a*_3_(*n*, **d**, *q*) is determined in a manner similar to above. That is, we set *q*_4_ = *q*_3_ + *f_D_*(**d**)*F_N_*(*n*)*m_d_p*_3_*n, q*_34_ to be the average of *q*_3_ and *q*_4_, and

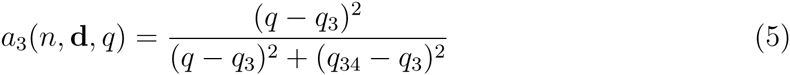

if *q* > *q*_3_ and 0 otherwise.

Note that the maximum fraction of neoblasts active in one unit of time is *p*_1_ + *p*_2_ + *p*_3_, which provides a link between the cell cycle and the physical meaning of the time unit (see **Parameters**).

### The Dynamics of Neoblasts

We write the dynamics as difference equations rather than differential equations for two reasons. First, even if written as differential equations, the dynamics have to be solved numerically, requiring conversion to difference equations. Second, the use of difference equations makes the balance for the cell dynamics more explicit. With these assumptions, the dynamics of neoblasts are

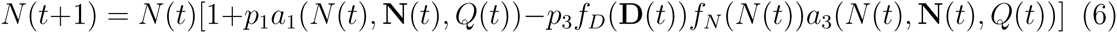

For those who prefer differential equations, one can proceed as follows. First replace *N*(*t* + 1) on the left hand side by *N*(*t* + Δ*t*) where Δ*t* is a suitably small unit of time. Second, define *r_i_* by *r_i_* = *p_i_*Δ*t.* Then we have

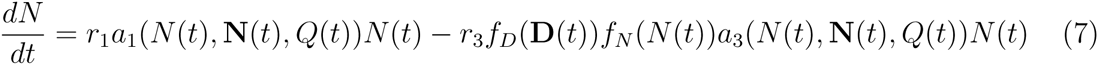

Thus, our difference equations are an Euler-method for the solution of the differential equations.

### The Production of Progeny

As described above, we envision an intermediate progenitor cell between neoblasts and fully differentiated cells. This progenitor is not mitotically active and may continue development to a fully differentiated cell or may return to the resource pool. This is an inefficient process, but important if food finding is stochastic and lineage commitment must be made before food is searched for. In light of the feedback functions described above, the total production of progenitor cells at time *t* is

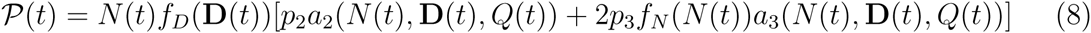

We assume that a fraction 1 – *f_Q_*(*Q*(*t*)) of these progeny are returned to the resource pool and that the remaining fraction complete differentiation. For computations we assume *f_Q_*(*q*) = 1 − *exp*(−*β_Q_q*), so that *f_Q_*(*q*) ≈ *β_Q_q* when *q* is small and *f_Q_(q*) → 1 as *q* increases. For cases in which we assume sufficient resources, we assume *f_Q_*(*q*) = 1.

### Dynamics of Differentiated Cells

To model the dynamics of differentiated cells, we must capture two processes: the mortality of differentiated cells and the distribution of progenitors across the diversity of differentiated cells.

We let 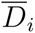 denote the target number of differentiated cells in the steady state planarian, which we assume is set by natural selection and thus exogenous to the dynamics of cells within the life of a planarian. Then 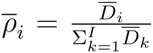 is the fraction of differentiated cells of type *i* in the steady state. Similarly, we let 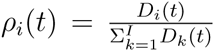 represent the fraction of differentiated cells of type *i* at time *t*. We assume that the rate of mortality of differentiated cells depends upon i) how far *D_i_*(*t*) is from 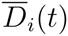, ii) how far *ρ_i_*(*t*) is from 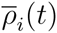, and iii) whether there are sufficient resources to maintain existing neoblasts and differentiated cells. In particular assume that the rate of mortality of differentiated cells of type *i*, given *N*(*t*) = *n*, **D**(*t*) = **d**, *Q*(*t*) = *q* is

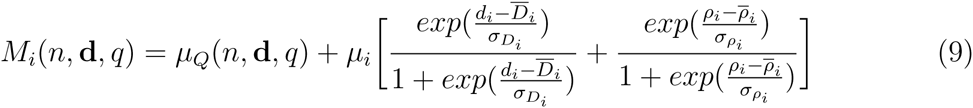

where *μ_Q_*(*n*, **d**, *q*) is the resource-dependent rate of mortality, which we determine as follows. We set *q_μ_* = 1.5*q*_4_ and assume that if *q* > 10*q_μ_* then *μ_Q_*(*n*, **d**, *q*) = 0 and otherwise

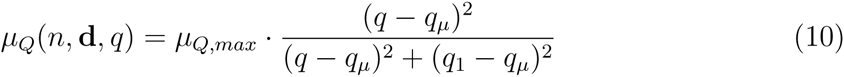

In the second term of Eqn 9 *μ_i_*, 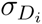 and 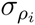 are fixed parameters. Note that when the planarian is in homeostasis with sufficient resources (*μ_Q_*(*n*, **d**, *q*) = 0), so that 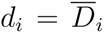 and 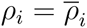, the second term on the right hand side of Eqn 9 is *μ_i_.*

To determine the allocation of progenitors across the different kinds of differentiated cells, we follow the relative need assumption as described above, the measure at time *t* for the need of the *i^th^* kind of differentiated cell is 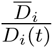 so that the relative need for the *i^th^* kind of cell is 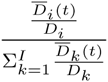. With these assumptions, the dynamics of differentiated cells are

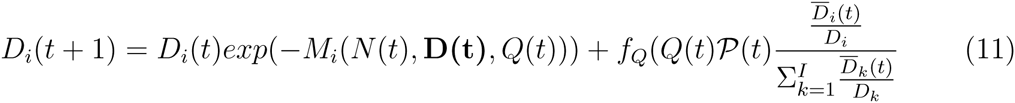

### The Dynamics of the Resource Pool

The resource pool increases by acquisition of resources from the environment, from progenitors that are directly returned to the pool and from differentiated cells that die. It decreases due to metabolism of neoblasts and differentiated cells and through cell divisions. We assume that resource gain from the external environment is *Y_e_D*_1_(*t*)^5^ where *Y_e_* is a metric of food availability in the external environment, *D*_1_(*t*) is the number of differentiated cells used for food gathering and *δ* < 1 is a parameter accounting for not all cells being able to accumulate resources from the external environment. Resources returned to the pool from a progenitor that dies are *γ_p_* and from a neoblast or differentiated cell that dies is *γ*. Since *m_r_* and *m_d_* denote the resource cost of metabolism and division, the dynamics of the resource pool are dynamics of the resource pool are

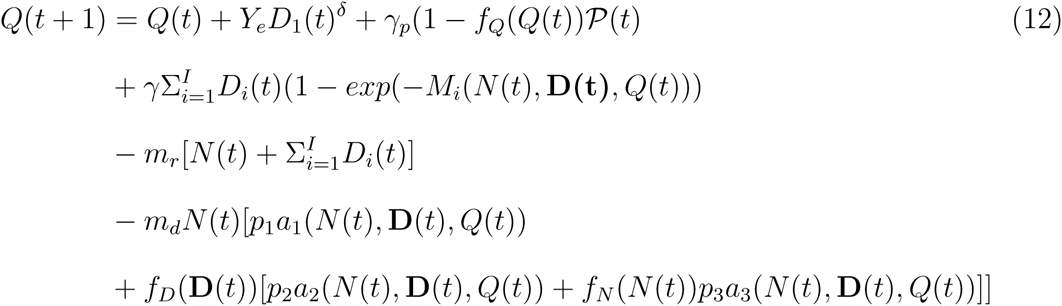

### The Steady State Under Sufficient Resources

Under sufficient resources, with overline denoting the steady state value of a dynamical variable, we assume 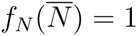, 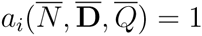 for all *i*, 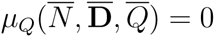, and 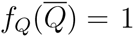. Thus none of the neoblast divisions are resource limited and all of the progenitors continue to full differentiation, rather than returning to the resource pool.

The neoblast dynamics (Eqn 6) become

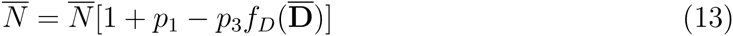

We assume that at the steady state all the feedback functions in Eqn 2 take the same value, so that for each *i*

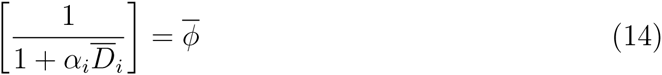

with 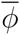 to be determined. Using Eqn 14 in Eqn 13 we have

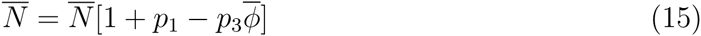

Note that this equation only makes sense if 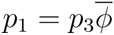. However, in light of Eqn 14, 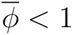, so we conclude *p*_1_ < *p*_3_ as a condition for the steady state.

In this steady state, the production of non-mitotic progeny is, from Eqn 8,

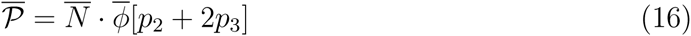

and Eqn 11 becomes

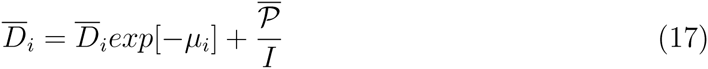

Using Eqn 16 in Eqn 17 we obtain

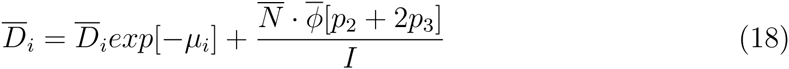

so that

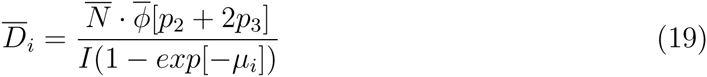

We now define

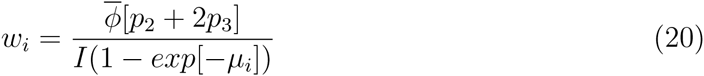

which allows us to write 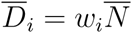. Consequently the fraction of neoblasts in the steady state is

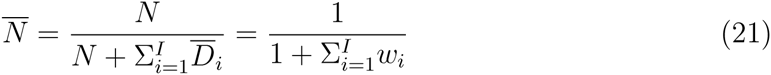

Thus we predict the same proportion of neoblasts in a planarian at the steady state, regardless of the number of cells and that this proportion is determined by the transition rates and rate of death of differentiated cells. In addition, the *μ_i_* will determine the relative abundance of differentiated cells in the steady state (and dynamically changing animal as well). For computations, we set *μ_i_* = *s_i_μ*_0_, where *μ*_0_ is the baseline rate of mortality for differentiated cells and *s_i_* is a modulator according to the kind of differentiated cell.

In the steady state, Eqn 12 becomes

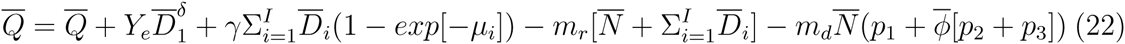

Since 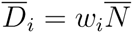 this becomes an equation for the number of neoblasts

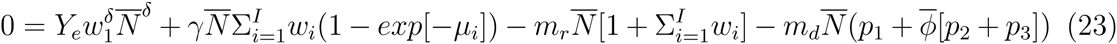

which we solve to obtain

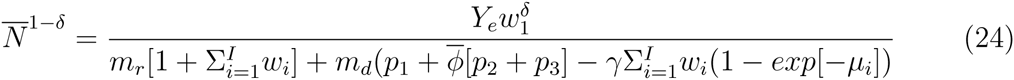

Once this equation is solved, we compute the number of differentiated cells from 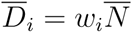 and then determine the *α_i_* by solving

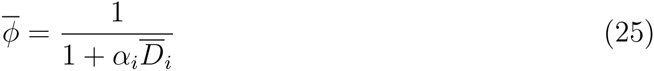

Since the steady state number of neoblasts, and thus of differentiated cells depends upon the level of food in the environment, environment-dependent feedback control is an emergent property of the model.

### Size-Cell Number Relationship

We compute the size of the planarian using the data from Table 1 of [15], which reports cell numbers and size for *Dugesia mediterranea* of 4, 7, 11, and 16 mm. We assume that size is determined only by the number of differentiated cells. Using those data, size *S*(*t*) at time *t* when the number of differentiated cells of type *i* is *D_i_*(*t*) is

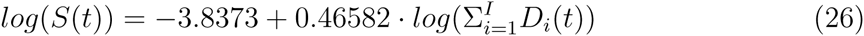

(*R^2^* = 0.99993 for the log-log plot).

### Growth Without Resource Constraints

If we assume that there are sufficient resources to support all divisions, then the full dynamics simplify considerably. That is, if we set *a_i_*(*N* (*t*), **D**(*t*), *Q*(*t*)) = 1, *μ_Q_*(*N* (*t*), **D**(*t*), *Q* (*t*) = 0, *f_Q_*(*Q*(*t*)) = 1, and the resource dependent mortality in Eqn 9 to 0, then

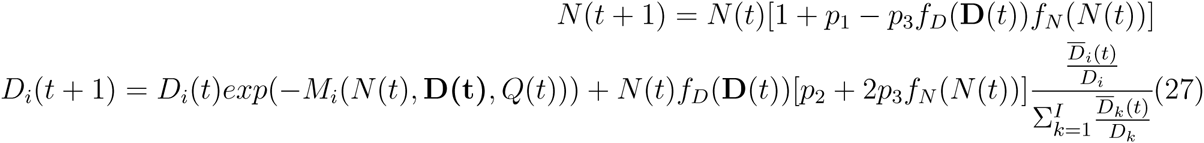

In this case, the dynamics of the resource pool are irrelevant. During such growth, the number of differentiated cells of type *i* dying in an interval of time will be *D_i_*(1 − *exp*(−*M_i_*(*N*(*t*), **D***_i_*(*t*), *Q*(*t*)) so that the per capita death rate of these differentiated cells is 1 − *exp*(−*M_i_*(*N*(*t*), **D***_i_*(*t*),*Q*(*t*)). This per-capita mortality is partitioned between the two types of non-steady state distribution of cells according to the relative values of the terms in Eqn 9. Our formulation allows us to separate mortality due to unbalance from the steady state level and mortality due to unbalance from the steady state proportions.

### A Simplified Version of the Model

For some questions, a version of the model with only one kind of differentiated cell suffices. We briefly describe that version now. We replace the mortality function in Eqn 9 by *μ_Q_*(*n*, *d, q*) + *μ* where *μ_Q_*(*n, d, q*) is interpreted as before and *μ* is a baseline rate of mortality. With this assumption the dynamics of neoblasts and the differentiated cell type become

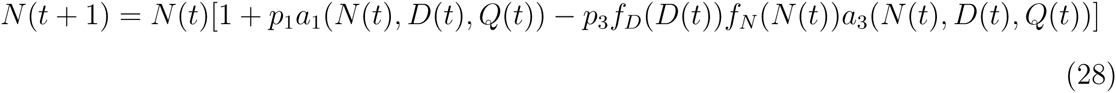

and

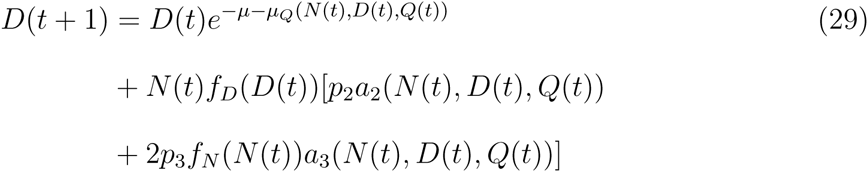

and the dynamics of the resource pool are

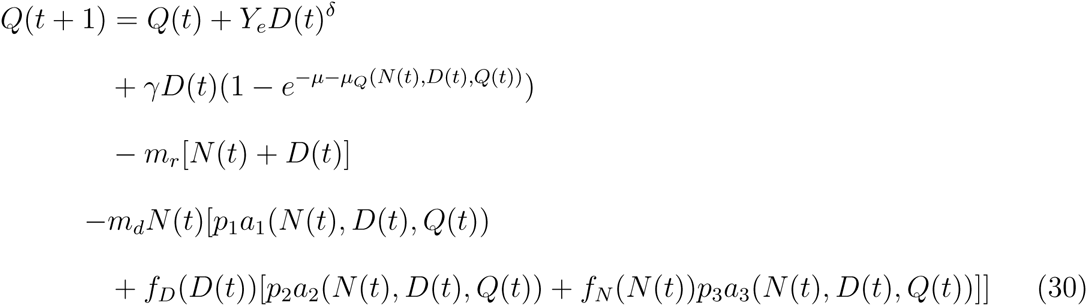

To compute the steady state without resource constraints, we set all the *a_i_*(*n, d, q*) = 1 and *f_N_*(*n*) = 1 and *μ_Q_*(*n, d, q*) = 0 (i.e., there are sufficient resources and neoblasts that transitions happen at their maximum possible value modified only by the feedback control from differentiated cells and there are sufficient resources that resource dependent cell death does not occur). Following the procedure as above for the full model, we find as before 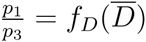 from which we conclude 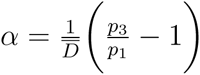 and

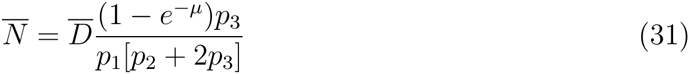

so that if we set 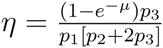 the fraction of neoblasts in this steady state is 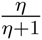.

In this simplified model, the steady state number of differentiated cells is determined from

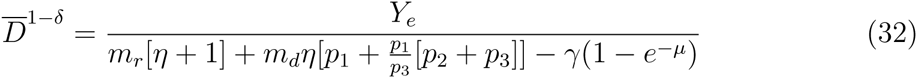

### Parameters

Although every parameter that is used in these models can be measured, most of them have not at this time. Indeed, one role of a paper such as this is to motivate empiricists to measure the parameters. We now explain how we determined the parameters. In general, we focus on the full model. When the simpler one differs from the full model, we explain the difference.

*Fundamental Transition Rates* Since we have used a discrete time model, in a unit of time that maximum fraction of neoblasts that are active is *p*_1_ + *p*_2_ + *p*_3_. This fraction implies a correspondence between a unit change in the time variable in the model and in physical time. The parameters are otherwise unconstrained except that *p*_1_ < *p*_3_ to ensure that a steady state exists. For computations here, we set *p*_1_ = 0.0001, *p*_2_ = 0.0005 and *p*_3_ = 0.00015.

*Food Gathering and Metabolic Rates* We assume that the exponent *δ* in Eqn 12 is described by the classic relationship between a linear variable and surface area, i.e. *δ* = 2/3 (qualitatively similar results are obtained with other choices, such as *δ* = 0.75). We choose metabolic rates in units so that the metabolic rate of a neoblast or differentiated cell is *m_r_* = 1.0 and assume that the cost of division is 4 times that, i.e. *m_d_* = 4.0. We assume that when a cell apoptoses 80% of its resources return to the resource pool so that *γ* = 0.8*m_d_*. In this framework, we understand food in the environment, *Y_e_* in Eqn 12, to be multiples of *m_r_*.

*Rates of Cell Death* We assume that the *μ_i_* in Eqn 9 are multipliers of a basic mortality rate *μ*_0_ so that *μ_i_* = *s_i_μ*_0_. We choose *μ*_0_ = .00015 and *s*_1_ = 0.75, *s*_2_ = 1.0 and *s*_3_ = 1. In the absence of resource constraints or deviations from homeostasis, the expected cell lifetime predicted from Eqn 9 is 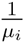, which is another way of setting the link between one unit of time in the model and chronological time. We set 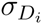 and 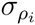 in Eqn 9 equal to 15% of the steady values of *D_i_* and *ρ_i_.* For the results reported in this paper, we assume sufficiently large resources so that *μ_Q_*(*n*, **d**, *q*) = 0.

*Feedback Controls* As described above, the parameters *α_i_* (or *α* in the simplified model) in Eqn 2 emerge from the steady state analysis. The choice of the functional form is somewhat arbitrary: we require that *f_D_* (**d**) declines as **d** increases and approaches 1 as **d** approaches 0 and the simple nonlinear form of Eqn 2 captures this idea without the risk of becoming negative as a linear function would; Taylor expanding these functions gives 1 − *α_i_D_i_.* Similarly, the choice of *q_ij_* and the exponent 2 used in the feedback control functions *a_i_*(*n*, **d**, *q*) in Eqns 1, 3, and 5 are arbitrary but capture the properties that we expect of such feedback functions. Finally, the feedback control *f_N_* (*n*) in Eqn 4 involves two parameters. We set the number of neoblasts at which the feedback control is 0.5 to 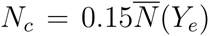 and the parameter characterizing the spread of this function 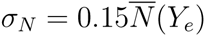.

### *In silico* Experiments

With the full model, we do the following. First, we compute the steady state size as a function of food level. From that we compute the strength of feedback control. We then compute the size, total mortality of differentiated cells, and fraction of neoblasts under a temporal pattern of food in which food is dropped and then subsequently increased but there are sufficient resources for all the *a_i_*(*n*, **d**, *q*) = 1 and *μ_Q_*(*n*, **d**, *q*) = 0. Fourth, we follow the dynamics of cells during remodeling following a division, again under the assumption of sufficient resources. To do, this we assume that the initial cell numbers are 50% neoblasts, and 10%, 30%, and 5% of the three kinds of differentiated cells respectively (rather than 25% neoblasts and 40%, 30%, 30% relative distribution of differentiated cells in the steady state).

Using the simplified model, we do the following. First, we follow the dynamics during growth and regrowth. Second, we consider an excision experiment: we grow a planarian and then at time t=70,000 the number of differentiated cells is reduced by 25%. For the x-ray experiment, we assume that at time t=70,000 the number of neoblasts is reduced by 25%.

## Competing Interests

The authors attest to no competing interests.

## Authors Contributions

MM developed initial forms of the models, which were then modified through discussions with AA and MB. MM did the computations. All authors contributed to the manuscript, with MM leading the writing.

## Acknowledgments

MM was supported by NSF grant EF-0924195; AA by BBSRC Research Grant BB/K007564/1 and MRC Research Grant MR/M000133/1.

